# A computational study on the role of parameters for identification of thyroid nodules by infrared images (and its comparison with real data)

**DOI:** 10.1101/2021.01.20.427415

**Authors:** José R. González Montero, Charbel Damião, Maira B. H. Moran, Cristina A. P. Fontes, Rubens Cruz Filho, Giovanna Balarini, Aura Conci

**Affiliations:** Institute of Computing, Fluminense Federal University, 24220-900, Niterói, Rio de Janeiro, Brazil; Department of Internal Medicine, Fluminense Federal University, 24033-900, Niterói, Rio de Janeiro, Brazil

## Abstract

According to experts and medical literature, a healthy thyroid gland, or a thyroid containing benign nodules, tend to be less inflamed and less active than one with malignant nodules. It seems to be a consensus that malignant nodules have more blood veins and it may be related to the maintenance of high and constant temperatures. Investigation of these characteristics, detectable by infrared sensors, and answering if they constitute patterns of malignancy are the aims of this work. Experiments considering biological heat transfer analysis by Finite Element numerical simulations are used to show the influence of nodule and patient characteristics on the identification of malignancy of thyroid nodule by thermography. The used and approved protocol for infrared examination are analyzed and simulated during all its phase that is on transient and steady state behavior, in order to verify how and when their influence can be really recognized in patients. Simulation results and the analysis of infrared exams show that the tissues between the skin and the thyroid, as well as the nodule size, have influence in superficial temperatures. Other thermal parameters of thyroid nodules are also investigated and show little influence on surface temperatures. These characteristics must be considered in nodular infrared analysis and diagnosis by thermography. The infrared examinations of patients that meet the hypotheses related to the vascularization of the nodule confirm the numerical results. All details of the physical parameters used in the simulations, characteristics of the nodules, and their complete thermal examinations are public and available, turning possible that the presented simulation could be compared with other types of heat transfer solutions. This study is a concrete contribution to the answer of under what conditions thermography can be useful in the identification of thyroid nodules.

**Author summary:** Thyroid nodules are very common health problems. These nodules may have different characteristics, and some of them could influence the temperature of the region. Many works in the medical literature report that the healthy thyroid and even benign nodules tend to be less inflamed and active than malignant nodules and therefore should exhibit some variation in patterns of behavior related to the temperature variation between them. The focus of this work is to analyze some parameters of the nodules and details of the patients that can influence the identification and diagnosis of thyroid nodules by infrared images. To reach the objective, simulations of bioheat transfer in the neck (using a simple neck geometry and Finite Elements Analysis in COMSOL Multiphysics Software) and real infrared examinations (performed with a FLIR Infrared Camera model SC620 and a proposed protocol) were analyzed. Our results show the thermal insulation effect of the neck fat tissue, and that the effect of nodule sizes can be decremented by a thicker layer of fat. The analysis of the nodule parameters as blood perfusion rate and metabolic heat, which could be related with the nodule vascularization (an important condition related with malignancy), suggest that the thermal effects of thyroid nodules on the neck surface are not sufficient to differentiate benign from malign, and for this other features or methods must be considered.

## Introduction

Although most thyroid nodules are benign, the possibility of malignancy should be considered. The importance of investigating thyroid nodules is related to the need of excluding the possibility of cancer, which occurs in 7% to 15% of cases depending on patient’s age, sex, family history, and radiation exposure, among other factors [1, 2].

Benign nodules are made up of similar mature cells, with characteristics similar to the healthy surrounding tissue [3]. These nodules present clear limits and are restricted to a single mass circumscribed by a capsule or adjacent tissues. They have a more slower and organized growth than malignant nodules, and are homogeneous in their morphology [1, 3].

Malignant nodules are characterized by a disordered and uncontrolled growth of cells [4], invasion of tissues, and spreading to other parts of the body, causing metastases [5, 6]. Cancerous cells divide themselves faster than normal ones and have infiltrating growth (enzymes that cause protein lysis and destruction of adjacent tissue) [7]. Cancerous cells are atypical, undifferentiated, immature, and functionally less specialized than their corresponding normal cells [8]. To grow and develop, such cells need nutrients, which causes the development of new blood vessels around them. This phenomenon increases vascularization and is named angiogenesis [9]. The nitric oxide, which is produced by cancer cells, interferes with the normal neural control of the blood vessel causing local inflammation and provides neoangiogenesis [10]. Central vascularization and a high rate of metabolic activity are important characteristics of malignant nodules [11]. Due to increased blood flow, malignant nodules could have higher and more constant temperature than the surrounding region and benign nodules; the verification (or contradiction) of this hypothesis is one of the main aspects of this study.

Infrared Thermography (IRT) uses camera and software to convert the infrared radiation emitted by objects into temperature values [12, 13]. Digital Infrared Thermal Imaging (DITI) [14] is a non-invasive technique based on IRT that allows us to visualize and quantify temperature changes in the skin surface [15]. DITI has been successfully explored in various fields of medicine, such as the determination of circulatory problems, assessment of the body’s reaction to a medication, physiotherapy treatments, study and diagnosis of various conditions, and detection of various cancers, especially breast tumors [15–19].

Few previous works had investigate the use of IRT to aid in the diagnosis of thyroid diseases. Some of them only analyze the thermal effects of hyperthyroidism and hypothyroidism by IRT [20, 21]. There are works on theoretical studies of the possibility of uses IRT in thyroid nodule diagnosis [18, 22–26], as well as paper evaluating the uses of IRT as complementary technique to help on the indication of biopsy of nodules suspected of malignancy [27]. However, there is no work on simulating a complete protocol of thermography specially including the transient phase: this aspect has been considered crucial on investigation the possibility of IRT for cancer diagnosis in previous work [28] and is accomplished in the present work.

The environmental, technical, intrinsic and extrinsic patient factors cited in [29] to use IRT for medical purposes were carefully considered in the simulated protocol of this work. This protocol is used to perform the infrared neck examinations of volunteers with benign and malign thyroid nodules on follow up by the medical doctors of our research group [30]. Even following this protocol for all patients, the analysis of preliminary thermographies ([31–33]) showed to be insufficient to promote a differentiate nodule diagnosis. These results could be related to the influence of more factors than those previously considered [29].

Clinical and biologically, the similarity or the differences of the nodules is evaluated by parameters completely diverse from those used in mechanical heat transfer analysis. Even the size, that is the most similar of all, has different ways of being measured: the length of the nodules in each direction, for instance, is measured by ultrasound exams [27] most of the time only by nodular higher and smaller diameters, without relation with the body axial planes or any fixed axial reference of the numerical model. Furthermore, thyroid nodules can be at different positions and depths inside the gland regardless of their diagnosis [27, 34]. Additionally, the vascular distribution of thyroid nodules is a very complex issue. In the literature is common to find reports of malignant nodules “poor on the number of veins” as well as “hypervascularized benign nodules” [35, 36]. Patients with thyroid nodules have different body constitutions, summarized by the relationship between body mass (weight) and height. Therefore, all these characteristics could produce different infrared thermal patterns even in patients with the same diagnosis for their thyroid nodules, suggesting that an internal heat source is dependent of the nodule echogenity and size [37]. Other variables related to the physical phenomenon of heat transfer are: the heat rates produced by nodules; the heat rates produced by organs and tissues inside the neck; the heat transfer mechanism from the inside to surface trough tissues; and the anatomical and physiological characteristics of patients and nodules. All of them must be considered jointly [29].

Based on the hypotheses that nodules with similar characteristics could produce similar thermal patterns, and when presenting different thermal patterns with the same features they could be related to nodule malignant or benign diagnostic, the objective of this work is to understand how parameters of thyroid nodules (blood perfusion rate, metabolic heat, size, etc.) and the patient characteristics influence the identification of thyroid nodules by infrared thermography. To reach this aim two studies are performed: The first does a bio-heat transfer analysis through numerical simulations by using the COMSOL Multiphysics® v5.2 [38] finite element software (license number: 1042008) in order to identify relevant elements and their relations. The second compares infrared examinations of volunteers with a definitive diagnostic with simulations to verify if the numerical founds are compatible with reality detected in some human beings. As far as we know both studies are done for the first time to support the diagnosis of this gland by thermography.

This work is part of the research “Acquisition, storage, and verification of the feasibility of the use of thermal images in the detection of thyroid diseases” (in Portuguese: “Aquisição, armazenamento e verifição da viabilidade do uso de imagens térmicas na detecção de doenças da tiróide”) designed to assess the importance of thermal imaging in diagnostic of thyroid nodules in accompanied patients at “Antônio Pedro University Hospital” (HUAP) of the Fluminense Federal University (UFF). The research was approved by the Research Ethics Committee of the UFF, it is registered with the number of Certificate of Presentation for Ethical Appreciation (CAAE): 57078516.8.0000.5243, and completely detailed in the public platform of the Health Ministry of Brazil (accessible at http://plataformabrasil.saude.gov.br) [30].

## Materials and methods

All details related to performed numerical simulations as well as infrared examinations of the volunteers are presented in the next subsections.

### Details of the bioheat numerical analysis

This section shows theoretical and practical details about the bioheat transfer analysis performed by using the COMSOL Multiphysics software [38]. Two types of numerical studies were accomplished to evaluate how and which parameters can influence the bioheat transfer related to a thyroid nodule: one considers the skin in thermal equilibrium with the environment, and the other introduces a forced ventilation in the beginning.

#### Neck simplified geometry

The 2D simplified geometry of the neck presented in Fig. 1-left was used for the basic model. This geometry is based on a consensus image (Fig. 1-right) of an average human neck. The consensus image corresponds to an axial view of the anterior part of the neck in the best position for thyroid representation on the transverse cutting plane.

**Fig 1.**
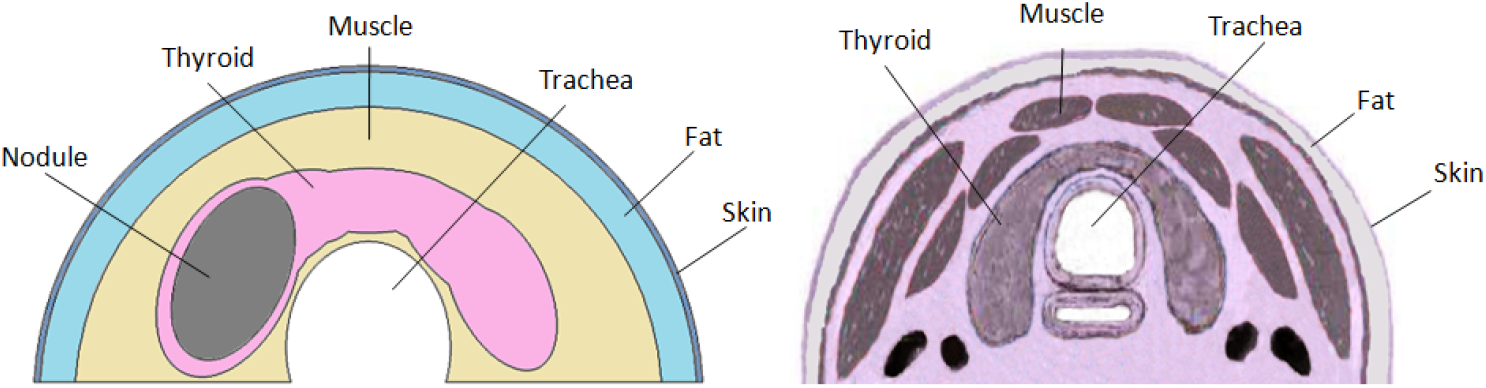
Simplified 2D geometry used (left) based on the averaged best view of the thyroid gland (right).

The simulated neck considers skin, fat and muscle layers, thyroid gland, and an elliptic nodule inside the thyroid lobe. The thicknesses of the skin and muscle tissues in Fig. 1-left are 0.1 *cm* and 1.0 *cm* [39], respectively. The thickness of the fat layer and the nodule size will be modified in the simulation to study their influence on the skin temperature.

Simple geometric elements (cylinders and ellipses) and logical operations (union and difference) available in COMSOL Multiphysics software were used to create this geometry [38].

The effects of the trachea, arteries, and jugular veins of the neck were disregarded. The simplified geometry was modeled by an extremely-fine triangular mesh using a proper tool of the used software, being its convergence tested in a preparatory stage of this research.

#### Mathematical and physical model

The Pennes’ bio-heat transfer model is used in the simulation [40]. This model considers the total energy balance and its storage, the internal energy rate, heat conduction, convection inside and outside the body and environment, as well as local heat generation [13]. At the same time, chemical and electrical effects are neglected in the model [37].

The temperature range is obtained for a homogeneous, solid, and linear biological medium with isotropic thermal properties [41]. The energy balance assumes that blood flow within the tissue is promoted at the capillary level, i.e., the capillaries are assumed to be oriented concerning their arterial and venous connections [41].

Pennes’ model uses the modified transient heat conduction equation and two heat sources, both per unit of time and volume: a heat source due to the metabolic effect, and a heat source due to the energy exchange between tissue and blood [41]. Besides, a source of external heat could be included but it does not match the protocol of exams considered in this work [42, 43].

Simulations considering no skin ventilation and superficial cooling and reheating were performed [28]. For the cooling period, a thermal perturbation over the neck surface with electric fan airflow was simulated [17, 30]. The reheating period is based on the return to thermal equilibrium with the environment (the examination room) immediately after the cooling period.

##### Formulation and conditions without skin ventilation

The mathematical formulation of the Pennes’ model at steady state is shown in Eq. 1. In this equation, *k* is the thermal conductivity, *w_b_* is the blood perfusion (blood flow rate per volume unit of tissue), *ρ_b_* is the blood mass per volume (or density), *c_b_* is the blood specific heat, *T_b_* is the arterial blood temperature at the capillary level, and *Q_m_* is the metabolic heat of the tissue.

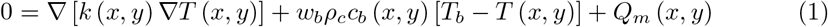

The boundary conditions are: the thermal convection (Eq. 2) [44] with the environment at skin surface (Γ_1_) using the convection coefficient *h* =10 *W*/(*m*^2^ *K*) (a typical value for natural convection) [22, 24, 26] and air temperature at *T_air_* = 25 °*C*, thermal insulation (Eq. 3) in the trachea surrounding region (Γ_2_), and prescribed temperature (Eq. 4) at *T_p_* = 37 ° *C* on the remaining surfaces (Γ_3_).

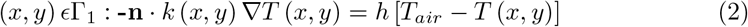

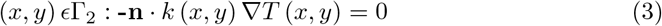

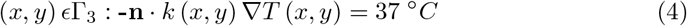

##### Formulation and conditions for cooling and reheating

Pennes’ model at transient steady is represented by Eq. 5. In this equation, *p* and *c* represent the specific mass and specific heat of tissue, respectively. The other parameters were already commented on previously.

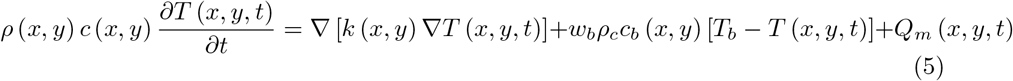

To simulate the thermal stress at the transient state due to the cooling with an electric fan airflow, the Eq. 2 representing the convection in the skin surface (Γ_1_), was also considered. However, in this case, a convection coefficient equal to *h* = 50 *W*/(*m*^2^*K*) was used to simulate forced thermal convection with the airflow [45]. The air temperature was maintained in *T_air_* =25 °*C* because the fan airflow hits the neck with pressure and speed, but with the same temperature. These values were selected because previous experiments performed allow getting temperatures very similar to the temperatures in the skin surface after five minutes of cooling in real examinations. Thermal insulation (Eq. 3) in the trachea surrounding region (Γ_2_), and prescribed temperature (Eq. 4) at *T_p_* = 37 °*C* on the remaining surfaces (Γ_3_), were also considered as boundary conditions. The solution obtained at the steady state was used as the initial condition for the cooling period. For this reason, each curve on the graphs in the session of “heat transfer analysis at transient state” begins differently related to their temperature.

To simulate the thermal reheating after the cooling period, the thermal convection (Eq. 2) in skin surface (Γ_1_) was considered again. In this case, the convection coefficient (*h* = 10 *W*/(*m*^2^*K*)) and air temperature (*T_air_* = 25 °*C*) were used to simulates the natural thermal convection with the environment. Thermal insulation (Eq. 3) in the trachea surrounding region (Γ_2_), and prescribed temperature (Eq. 4) *T_p_* = 37 °*C* on the remaining surfaces (Γ_3_), were also considered. The last solution obtained in the cooling period was used as the initial condition for the reheating period.

#### Thermophysical parameters used in experimentation

Table 1 presents the values of thermophysical parameters for every simulated tissue [22, 24, 46]. The *ρ_b_* and *c_b_* parameters are considered to be equal to specific mass (*ρ*) and specific heat (*c*) of the tissues.

**Table 1.**
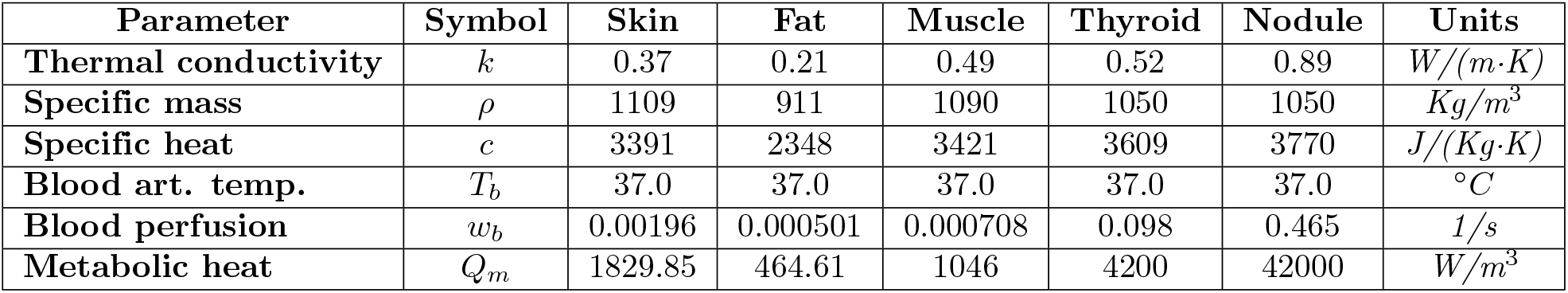
Used tissues’ thermophysical parameters.

| Parameter | Symbol | Skin | Fat | Muscle | Thyroid | Nodule | Units |
| --- | --- | --- | --- | --- | --- | --- | --- |
| Thermal conductivity | $k$ | 0.37 | 0.21 | 0.49 | 0.52 | 0.89 | $W/(m \cdot K)$ |
| Specific mass | $\rho$ | 1109 | 911 | 1090 | 1050 | 1050 | $Kg/m^3$ |
| Specific heat | $c$ | 3391 | 2348 | 3421 | 3609 | 3770 | $J/(Kg \cdot K)$ |
| Blood art. temp. | $T_b$ | 37.0 | 37.0 | 37.0 | 37.0 | 37.0 | $^\circ C$ |
| Blood perfusion | $w_b$ | 0.00196 | 0.000501 | 0.000708 | 0.098 | 0.465 | $1/s$ |
| Metabolic heat | $Q_m$ | 1829.85 | 464.61 | 1046 | 4200 | 42000 | $W/m^3$ |

#### Details of steady state experimentation

Four aspects are considered in the steady state experiments: two are size related and two representing heat transfer parameters.

The size related considers nodule and fat sizes. The fat tissue layer varies among individuals. Fat thickness is associated with the body mass index (BMI) and can be measured by ultrasound (US) examinations [35, 36]. Three sizes of nodules as shown in Fig. 2 and four fat layer thickness (0, 0.3, 0.6, and 1.2 *cm*) were considered (as shown in Fig. 3).

**Fig 2.**
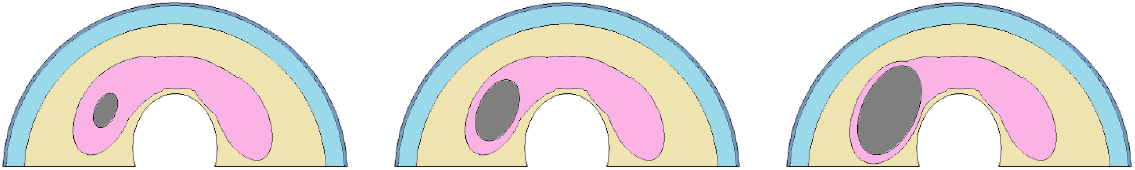
Variations of nodule sizes: (0.7; 1.2 *cm*) Small, (2.2; 1.26) *cm*) Medium, and (3.14; 1.80 *cm*) Large.

**Fig 3.**
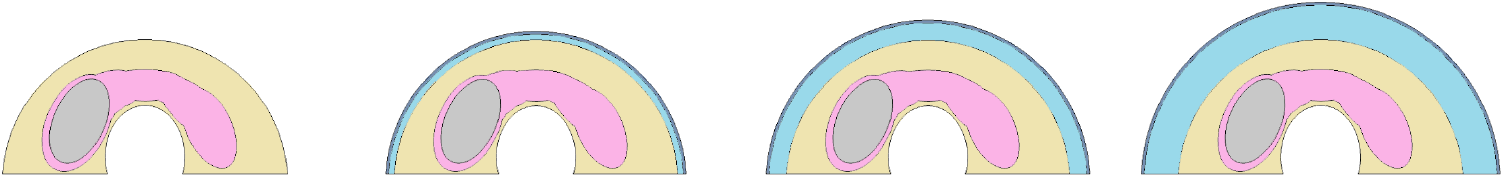
Variations of fat tissue thickness from 0 *cm* (first), 0.3 *cm* (second), 0.6 *cm* (third), to 1.2 *cm* (fourth).

As mentioned before, the metabolic heat (*Q_m_*) and blood perfusion rate (*w_b_*) of a thyroid nodule are vague parameters. Variations of these characteristics must be consistent with benign or malignant types of nodules, because they are the parameters that can be differentiated between behaviors compatible with vascularization and heat production, such as described by some works in the medical area [47, 48]. Table 3 shows the possible values of metabolic heat and blood perfusion rate that were used. The lower values of each parameter in this table are that of normal thyroid tissue, and the other two are the same used in [24].

#### Details of experimentation at transient state

As commented before, to the simulate the examination protocol used in the patient acquisitions, all studies at transient state were done in three phases: 1) the problem at steady state was resolved; 2) a cooling period during a specific time is simulated using the solution of the first phase as the initial condition, and 3) the reheating period is also simulated during a specific time using the last solution of the cooling period as the initial condition.

Each possible element that can present influences on the heat transfer from the nodule to the skin surface were evaluated by the same software used previously. The four fat tissue thickness (0, 0.3, 0.6, and 1.2 *cm*), the variations of nodule diameters from (0.7; 1.2 *cm*) Small, (2.2; 1.26 *cm*) Medium, to (3.14; 1.80 *cm*) Large, the metabolic heat and blood perfusion for the nodule used in the simulations at steady state were also inspected now. The same parametric configuration used in the simulations at steady state were used in the transient state.

Besides, a fixed point was located in the skin surface in front of the nodule, in the position of the red circle in Fig. 5-left for each of these four fat tissue thicknesses. From each point, temperature series were constructed with temperature values taken at every 15 seconds during the cooling and reheating periods. In some cases, symmetrical points of this concerning to the body axial axis are investigated, as well. Temperature series were also obtained from a fixed point located in the skin surface in the front of the nodule, in each simulation.

### Details for parameter analysis using infrared examinations of the volunteers

Data from eight volunteers, patients of the (HUAP/UFF) University Hospital [49], were selected to verified and compare real data with the numerical findings, and to show the influence of the nodules and patients parameters. Solid thyroid nodules were encountered in each of them and they were examined by using the same image acquisition protocol [30].

The used protocol establishes orientations in all levels (i.e. from patient’s orientation to technical and environmental requirements) and the methodology for the exam based on Dynamic Infrared Thermography (DIT) to evaluate the temperatures on the neck surface during the return to thermal equilibrium after thermal stress.

The thermal stress consists of neck cooling by an airflow (made by an electric fan) until it reaches a mean value of 30 °*C* or for 5 minutes, as maximum cooling time [42] [43]. After thermal stress, 20 infrared images are acquired, every 15 seconds for 5 minutes. Additionally, more one image is captured to indicate the position of the nodules with an insulating thermal material (that is fixed in the patients by their accompanying medical doctor after the identification of nodular position by touch or based on previous ultrasound examinations).

All infrared examinations were performed using an FLIR infrared camera model SC620. The camera sensor has NETD (Noise Equivalent Temperature Difference) value less than 40 *mK*, capture range of −40 °*C* to 2000 °*C*, and produce thermograms with dimensions of 640 ×480 pixels.

Figure 4 shows the last thermogram of the infrared examinations from of the eight patients, where the region of nodules was marked with a thin blue line in each thermogram (this line was done by one of the medical doctor coauthors of this work, that responsible for the patient and, the nodule position was confirmed by the others medical doctors based on previous exams).

**Fig 4.**
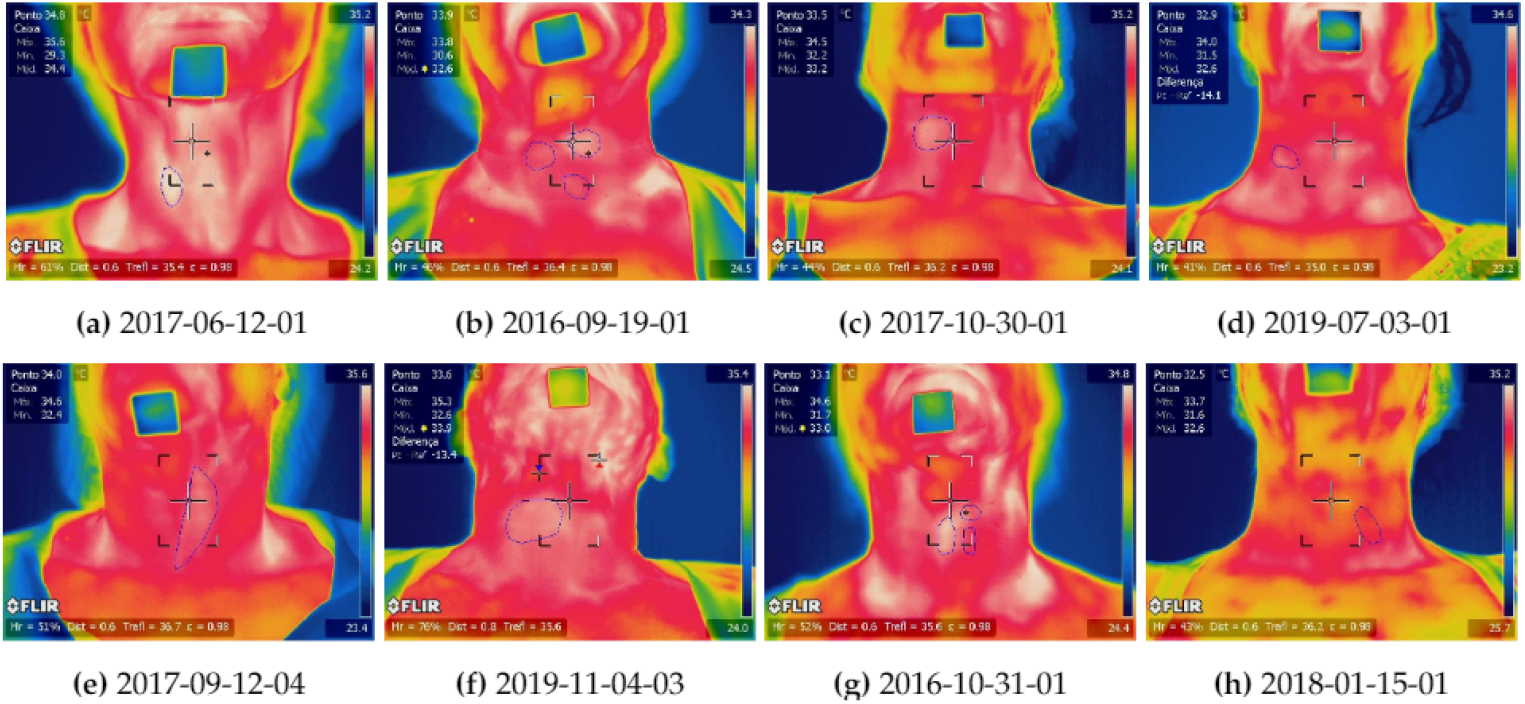
The last thermogram of infrared examinations of patients, where is possible to see the nodule position identified by endocrinologists.

Table 2 condenses all data: the dimensions, the principal ultrasound characteristics (presence of micro-calcifications, complete halo, regular contours, heterogeneity, and Chammas vascularization pattern), the cytopathological report (Bethesda classification), the body temperature related to these patients, and the confirmed (cytologically) diagnostic of the nodule (B - benign and M - malignant).

**Table 2.**
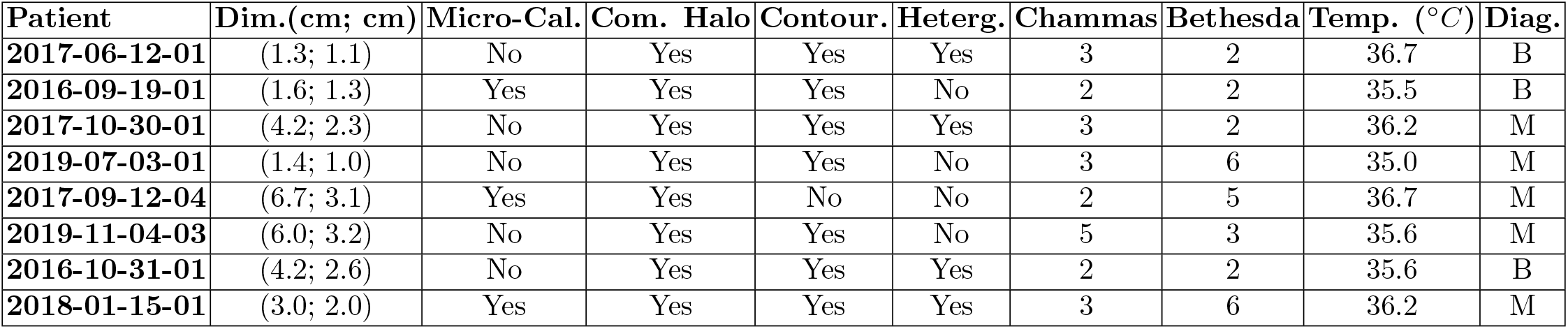
Dimensions, ultrasound characteristics, cytopathological result, and diagnosis of the nodules of the volunteers.

#### Influence of fat tissue thickness

To show the influence of fat tissue thickness using real data, two volunteers were selected. They are the patients identified as 2017-06-12-01 and 2016-09-19-01 in the database and can be seen in Fig. 4-a and Fig. 4-b.

The nodules of both patients have similar dimensions, the same pattern of vascularization (the third level in the Chammas scale [50]), and the same risk of malignancy in the cytopathological examination (6), as well. However, these patients present diverse BMI: 2017-06-12-01 has a thin layer of fat tissue on the neck, while the patient 2016-09-19-01 has more fat tissue.

#### Influence of nodule size

To show the influence of nodule size using real cases, the patients 2017-10-30-01 (Fig. 4-c) and 2019-07-03-01 (Fig. 4-d) of the database were selected. Their nodules have similar ultrasound characteristics and pattern of vascularization, but different dimensions, i.e., the nodule dimensions of these patients are (4.2; 2.3 *cm*) and (1.4; 1.0 *cm*), respectively.

#### Influence of vascularization

To compare the influence of vascularization of the real data and simulations two pairs of patients were selected. The first pair is composed by patients 2017-09-12-04 (Fig. 4-e) and 2019-11-04-03 (Fig. 4-f) in the database. Its nodules have similar dimensions and ultrasound characteristics, but different pattern of vascularization (in the Chammas scale).

The second pair (2016-10-31-01 (Fig. 4-g) and 2018-01-15-01 (Fig. 4-h)) presents nodules with different diagnostic. The patient 2016-10-31-01 has three benignant thyroid nodules, but only the biggest (the first from left to right in Fig. 4-g) was considered. The patient 2018-01-15-01 has a malignant thyroid nodule. These nodules have similar dimensions and ultrasound characteristics, as shown in Table 2. Both patients were also selected because they are skinny (i. e. similar body mass index), decreasing a possible influence of fat tissue. Moreover, their nodules have similar peripheral vascularization (second and third levels in the Chammas scale [50]).

## Results

The thermal energy transfer occurs from a high temperature place to a lower temperature location and may occur under steady or unsteady state conditions. Under steady state conditions, the temperature within the system does not change with time. When the temperature is time related, the system is considered under unsteady state conditions. Both states are analyzed in the experiments of this work.

First, it is important to mention that for all numerical studies carried out, a mesh convergence analysis was performed to verify the adequacy of the discrete model used on the representation of the continuous part of the human body modeled: the neck region. In the used mesh, the thyroid nodules are considered ellipsoids, and a simplified geometry is used to represent the elements of the body in the neck region under analysis (presented in *Neck simplified geometry* subsection).

Additionally, data of eight volunteers (selected from patients on follow up in the university hospital considering cases that exams and data better match the simulated hypothesis of the numerical study) are used to demonstrate how much the simulation match the reality of infrared examinations by using the same protocol.

### Bio-heat transfer analysis at steady state

This subsection presents the results of simulating the bio-heat transfer when the object under study is in equilibrium with the environment. Additionally, it shows how the body anatomy and tumor characteristics (specifically the nodule size and fat tissue thickness) could influence the temperature transmission up to the skin.

#### Variation of temperatures from inside the body to the neck surface

Figure 5-left shows the temperature variations represented by the color scale in the right (going from navy-blue = 34.0 °*C* to dark-red = 38.5 °*C*), computed by one of the several numerical simulations done when an elliptic nodule (with diameters of 3.14 and 1.80 *cm*) is considered inside a thyroid (in a neck with fat tissue layer equals to the human average value, which is of 0.6 *cm* of thickness for the frontal region). There are two lines and four points that must be noted in Figure 5-left. The lines are those labeled “cut line” on Figure 5-left horizontal axis. The red one crosses the nodule and the black is symmetrical to the red line considering the middle of the image. The point represented by the green triangle is the nodule center. The red circle points the beginning of the red line, its location on the skin and zero horizontal coordinates of the Figure 5-right. The black square and cyan diamond are symmetrical to the green triangle and red circle respectively. The red numbers in the perimeter of Figure 5-left represent the distance of the neck position surface tom the symmetrical axis (in the front of the neck). In other words, the red numbers in Fig. 5-left indicate the length of the arc at the skin surface (for a neck with a perimeter of 36 *cm* approximately) when a thickness equals to 0.6 *cm* is considered for the fat tissue layer.

**Fig 5.**
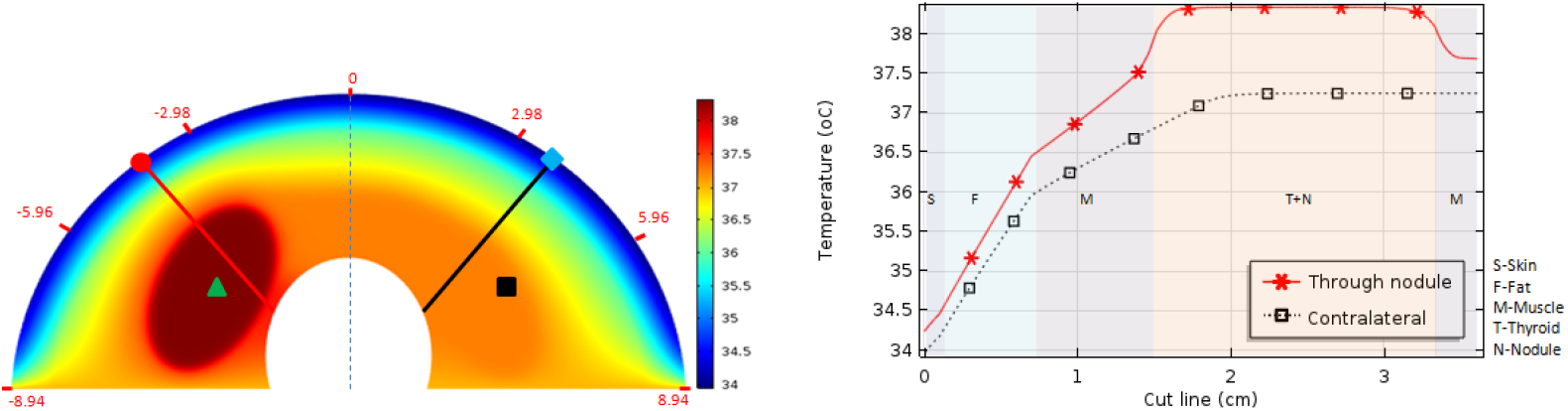
Temperature variations inside the neck represented in colors (left) and graphically along symmetrical lines (right). They are computed by finite element simulation considering a big elliptic nodule (17.76 *cm*^2^) and a medium fat tissue layer (thickness of 0.6 *cm*). Both (red and black) lines represent the temperature going from the surface to the trachea: they across the skin (S), fat (F), muscle (M), and thyroid (T) layers represented by the smooth colors. The red line goes through the nodule (N) as well.

The temperature variations presented in Fig. 5-right, from the skin to the trachea, are computed by the numerical simulation along contralateral lines considering the sagittal plane. The red line crosses the nodule and the black line (symmetrically positioned to the red one) crosses only the healthy thyroid lobe. As can be seen, the temperature variation into the fat layer for both lines was approximately equal to Δ*T_F_* = 2.0 °*C* showing the great importance of the fat tissue on the isolation of the internal body temperature from the superficial temperature.

It is interesting to note that temperatures of approximately 37.2 °*C* were obtained in the right thyroid lobe region (black square position), which is the region without a nodule corresponding to normal internal body temperature. Temperatures around 38.3 °*C* were found in the central region of the nodule (in the green triangle position).

The simulation output temperatures of approximately 34.3 °*C* in the neck surface related to the thyroid nodule region (around the red circle), and 34.0 °*C* in the contralateral region, that is the symmetric point in respect to the dashed line in the figure that represents the human sagittal plane (the point marked with a cyan diamond). This indicates a temperature variation of approximately Δ*T_CL_* = 0.3 °*C* in the neck surface caused by the presence of the nodule when compared with the healthy region (for this specified nodule size and fat thickness).

#### Influence of nodule size and fat tissue thickness in the surface temperature

There are four graphs in Fig. 6, each one with three curves showing the thermal distribution on the skin surface of the neck obtained from the simulations of nodules with three sizes (black = Large, red = Medium, and blue = Small). In the graphs four fat tissue thickness: 0 *cm* (Fig. 6-top-left), 0.3 *cm* (Fig. 6-top-right), 0.6 *cm* (Fig. 6-bottom-left), and 1.2 *cm* (Fig. 6-bottom-right) are considered. For these graphs: the central position (0 *cm* in the horizontal axis) is related to the dashed line in Fig. 5-left that represents the human sagittal plane; the negative and positive values in the horizontal axis are related to the left and right part from the dashed line, respectively. In these simulation, the parameters used for metabolic heat is *Q_m_* = 42,000 *W/m*^3^ and for the blood perfusion rate is *w_b_* = 0.465 1/*s* [24]. These values can be supposed representative of malignant nodules.

**Fig 6.**
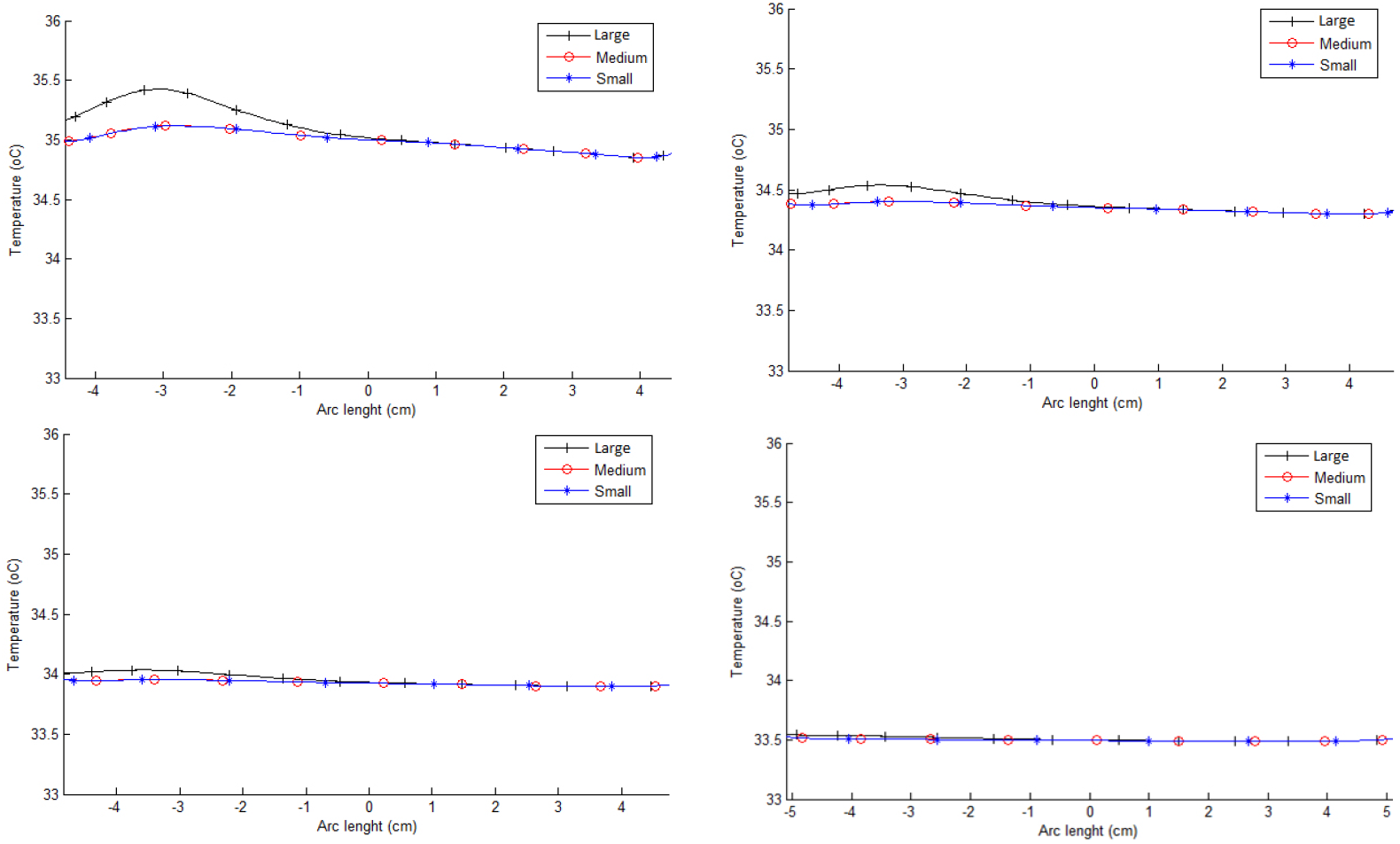
Superficial temperatures for Large (black line), Medium (red line) and Small (blue line) nodules of necks with fat tissue layer thickness from 0 *cm* (top-left), 0.3 *cm* (top-right), 0.6 *cm* (bottom-left), to 1.2 *cm* (bottom-right).

As can be seen, the thermal influence of the nodules on the skin surface decreases as the thickness of the fat layer increases. There is a perceptible surface temperature variation between the point just in front of the nodule (around −3 *cm* of the horizontal axis of the graphs, that represent the neck superficial position) and its contralateral point (around 3 *cm* of the horizontal axis of the graphs) for the simulation using the thinnest fat layer (Fig. 6-top-left). However, this variation (between symmetric points) is almost imperceptibly when the fat thickness increases, as can be seen in the other graphs of this Fig. 6. There is not a nodule significant effect for the fat layer with 1.2 *cm* thickness. Moreover, temperature variation between symmetrical superficial points is proportional to the size of the nodules (i.e. increase when the area of the nodules increases). Note that the fat layer can change the superficial temperature maximally 2 °*C* and the nodule size 0.6 °*C* (for the same neck). Therefore, the simulations presented in Fig. 6 show that the thermal insulation effect of the fat tissue layer is much more important than the nodule size, although both present the expected influence on surface temperature.

#### Influence of the metabolic heat and the blood perfusion rate in the neck heat transfer

The metabolic heat (*Q_m_*) and blood perfusion rate (*w_b_*) of thyroid nodules are very complex parameters of thermal sciences to be quantified. They are very used in thermal engineering and heat transfer; their values must be associated with benign or malignant types of nodules because they are the unique parameters that can distinguish the behaviors related to vascularization and heat production described (as commented before) in biomedical works [47, 48].

In case of a malignant thyroid nodule, Bittencourt et. al. [24] adopt for these parameters, values of *Q_m_* = 42,000 *W/m*^3^ and *w_b_* = 0.465 1/*s* using other malignant tumors as reference.

For normal thyroid tissue, the values of these parameters can be found as *Q_m_* = 4, 200 *W/m*^3^ and *w_b_* =0.098 1/*s* [22, 24, 46].

We did not find reports that mention these features for benign nodules up now. However, here we present the hypothesis that it makes sense to consider that these values for benign nodules would be intermediate, or some combination between these extreme values. In other words the values for normal thyroid tissue and malignant thyroid nodule can be considered as upper or lower bounds of possible values of these parameters.

Table 3 shows the four possible combinations from values of metabolic heat and blood perfusion rate obtained from the literature for normal thyroid (second row) and malignant tumor tissues (fifth row of the table). Curves in Fig. 7 shows the thermal distribution on the neck surface computed considering all combinations of the values of metabolic heat and blood perfusion for the thyroid nodule. In these simulations, the best possible conditions of the other parameters for skin temperature difference observation between contralateral points were used (that is, the biggest nodule with diameters of 3.14 and 1.80 *cm* and smaller possible fat tissue thickness: 0 *cm*). Although it is very difficult to find a correct relation among heat transfer parameters and biomedical behavior, the simulations presented in Fig. 7 shows that for any combination that could be representative of malignant or benign nodules, it is not possible to detect different behavior among these curves. All combination promoted the same modification on the skin’s surface between contralateral positions of 0.5 °*C*.

**Fig 7.**
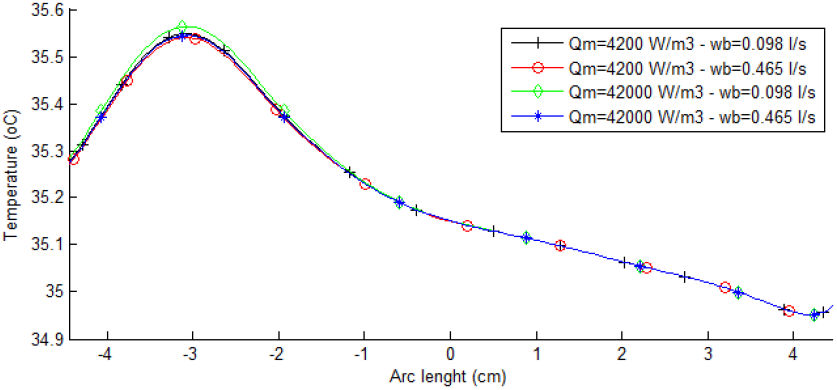
Surface temperatures obtained from the four possible combinations of metabolic heat and blood perfusion values considering no fat tissue and the biggest thyroid nodule.

**Table 3.** All combinations of metabolic heat and blood perfusion for thyroid nodule used in the simulations.

| Metabolic heat ($\text{W/m}^3$) | Blood perfusion ($\text{1/s}$) |
| --- | --- |
| 4,200 [22,24,46] | 0.098 [22,24,46] |
| 4,200 [22,24,46] | 0.465 [24] |
| 42,000 [24] | 0.098 [22,24,46] |
| 42,000 [24] | 0.465 [24] |

### Bioheat transfer analysis at transient state

The same previous aspects are from now analysed under transient conditions. Normally, the transient state is a precursor of steady state because certain time must pass after heat transfer be initiated, for a system reaches a steady state condition. Moreover, no system remains under a transient condition forever: the energy transferred into it changes. So the energy within the system presents alteration up the equilibrium with consequent temperature modifications.

This subsection shows the results of the performed simulations at the transient state. In the patient’s examinations reported in the next subsection such a condition is induced. Briefly, the simulations at the transient state here attempt to reproduce the infrared exam of the thyroid performed in our university hospital to be discussed in the next subsection. There, in the infrared exam of the thyroid, the neck surface is cooled with an airflow until reaching 30 °*C* or during five minutes under 25 °*C* (that is the room temperature). Then, the airflow is suspended and an infrared thermogram is captured every fifteen seconds during five minutes, in the reheating (recovery or warm) process [42, 43].

In the simulations, the cooling process is made up using air with a convection coefficient of *h* = 50 *W*/(*m*^2^*K*) and 25 °*C* for room temperature (the same of the infrared exam). The final configuration of the cooling process is used as the initial condition to the reheating process, then the reheating stage is simulated using natural ventilation with convection coefficient *h* =10 *W*/(*m*^2^*K*) and room temperature of 25 ° *C*. The simulation outputs every fifteen seconds are used to construct the temperature series presented.

#### Influence of fat tissue thickness and nodule size

The four curves in the graph of Fig. 8-left show the temperature series related to the red circle at the beginning of the red line in Fig. 5-left, that is located in the skin surface, computed by the four fat tissue thickness considered (0, 3, 6, and 12 *cm*). These results were obtained using the biggest nodule (with diameters of 3.14 and 1.80 *cm* or area of 17.76 *cm*^2^), the metabolic heat of *Q_m_* = 42,000 *W/m*^3^, and blood perfusion rate of *w_b_* = 0.465 1/*s* for the nodule. In the same way as in the steady state, the lower temperature variations (curve with black-triangle details) were obtained in the skin surface when the major fat tissue thickness was considered, showing that this layer reduces the influence of internal conditions in the neck surface temperature.

**Fig 8.**
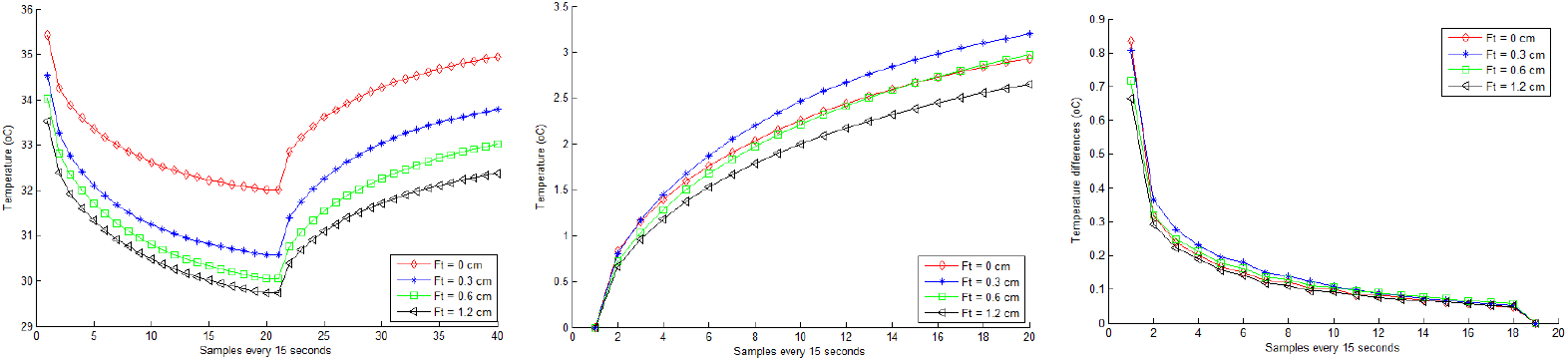
Influence of fat tissue thickness on the skin temperature just in front of the nodule (the red point in Fig. 5) over the time during all the exam simulated protocol (left graphic); temperatures at same position on natural reheating stage (center graphic) but considering each curve starting with the same temperature for better comparison; and the rate of superficial temperature at same point, i.e. differences between two consecutive temperatures (right graphic).

Fig. 8-left shows that the temperature limits for the curve representing the biggest fat tissue (1.2 *cm*) are: 33.6, 29.7, and 32.4 °*C*. These limits for the opposite case, the numeric abstraction representing a tendency of the fat layer to be on the lower possible case, i.e. having no fat tissue (0 *cm*), are 35.5, 32.3, and 35.3 °*C*. For the others fat thickness (i.e. between these values) the curves present limits almost visually proportional. Such variations are around 4 °*C* in cooling and 3 °*C* on warming. Note that they are greater than to any other variation in the steady state (even due to the size of the nodule) showing the importance of inducing a dynamic examination in this area of the human body for all types of patients under study. The thermal insulation is proportional to the thickness of the fat layer.

The curves in Fig. 8-center show only the reheating period of the previous temperature series (due to the infrared examinations of patients be performed in the reheating period) and related to its first value (i.e. they show the temperature series starting with the same value in order to promote a better form of representation the total variation of each one). As can be seen, temperatures related to the fat tissue layer of 1.2 *cm* (curve with black-triangle details) are smaller than those related to the fat tissue layer of 0.6 *cm* (curve with green-square details), and such temperatures are smaller than those related to the fat tissue layer of 0.3 *cm* (curve with blue-plus details) as well. The position of the null thickness curve (red one) is not representative, because this supposition of no fat layer must be seen as a tendency and not something real (in itself). In other words, its position in this graph may be a numerical inconsistency related to the abstraction realized by such an initial premise.

The curves in Fig. 8-right show the difference between two consecutive temperatures (Δ_*j,i*_ = *S_j,i_* – *S_j,i–1_*) in each of the temperature series (*S_j_*) starting in the reheating period. As the difference between each respective time is constant these curves can be directly associated to the rate of temperature changes. The higher heating rate appears for the thinner layer of fat (curve with blue-plus details related to 0.3 *cm*). Comparing the other two thicker layers: the speed of temperature change is proportional to the reduction of the fat layer. Or the speed of temperature change is inversely proportional to the thickness of the fat layer. (Again the relative position of the curve related to the null thickness is not representative).

The numerical experimentation performed allow to state that: (1) the simulations of the four fat tissue thicknesses at the transient state show the same insulation thermal effect of them in the steady state; (2) induced cooling and its relaxation can be measured in the neck surface for its temperature changes for any fat tissue thickness; and (3) any influence of the heat produced by a thyroid nodule on the surface of the skin will be mitigated if the layer of fatty tissue is large.

The three curves in the graph of Fig. 9-left show the temperature series related to the same point (marked by the red circle in Fig. 5-left) located in the skin surface considering three nodule sizes: Large (diameters of 3.14 and 1.80 *cm*, area = 17.76 *cm*^2^); Medium (diameters: 2.2; 1.26 *cm*, area =8.71 *cm*^2^); and Small (diameters: 1.2; 0.7 *cm*, area =2.64 *cm*^2^).

**Fig 9.**
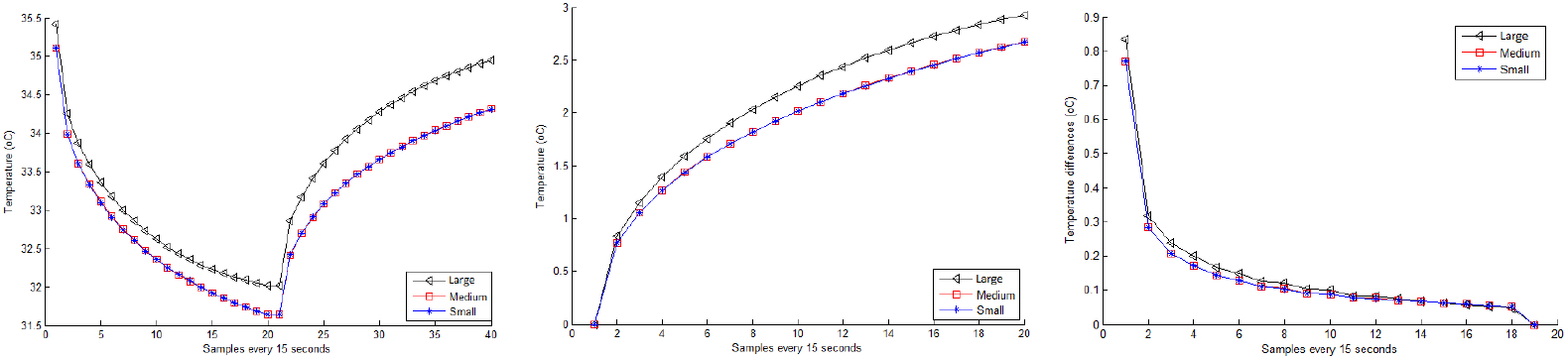
Influence of nodule area in the superficial temperature in front of the nodule over the time during the examination simulated protocol (left graphic); in the natural reheating stage (center graphic) considering each curve stating with the same temperature for better comparison and influence of nodule area in the velocity of superficial temperature change, computed from differences between two consecutive temperatures (right graphic).

The curves in Fig. 9-center also show the reheating period of these three temperature series related to the first value of each of them. These results were obtained using the smaller possible fat layer (fat thickness = 0 *cm*), metabolic heat of *Q_m_* = 42, 000 *W/m*^3^ and blood perfusion rate of *w_b_* = 0.465 1/*s* for the nodules. They show that the larger nodule presents the highest temperature (curve with black-triangle details) variations. No differences were found between the temperatures of the two smaller nodules. This analysis shows that larger nodules present more influence on the skin surface in the transient examination as occurs in the steady state study.

Figure 9-right show the temperature differences between two consecutive temperature values (Δ_*j,i*_ = *S_j,i_* – *S_j,i−1_*) in each of these three temperature series (*S_j_*) starting in the reheating period. The curves show that the nodules with the same parameters and different sizes did not show important differences in the rate of heating on the skin surface.

It is important to note that the final temperature in the reheating stage (in all the graphs of these studies) are less than the initial temperatures at the start of the cooling stage for all fat tissue thickness and nodule sizes considered. This shows that five minutes for the reheating stage is not sufficient to reach the same initial temperatures. The temperature before the cooling stage will only be reached in more time than that suggested by the acquisition protocol.

#### Influence of metabolic heat and blood perfusion rate

Temperature series in Fig. 10 were performed considering the larger nodule (diameters equal to 3.14 and 1.80 *cm*) and no fat layer (i.e. a thickness of *F_t_* = 0 *cm*). The three graphs were obtained considering all combinations of metabolic heat and blood perfusion (showed in Table 3). As can be seen, no differences were obtained for any combinations of these parameters for the nodule. This means that whatever could be the combination of them, within a range found in the literature, that represent a malignant or benign behavior, it is impossible to detect skin surface difference on temperature related to them.

**Fig 10.**
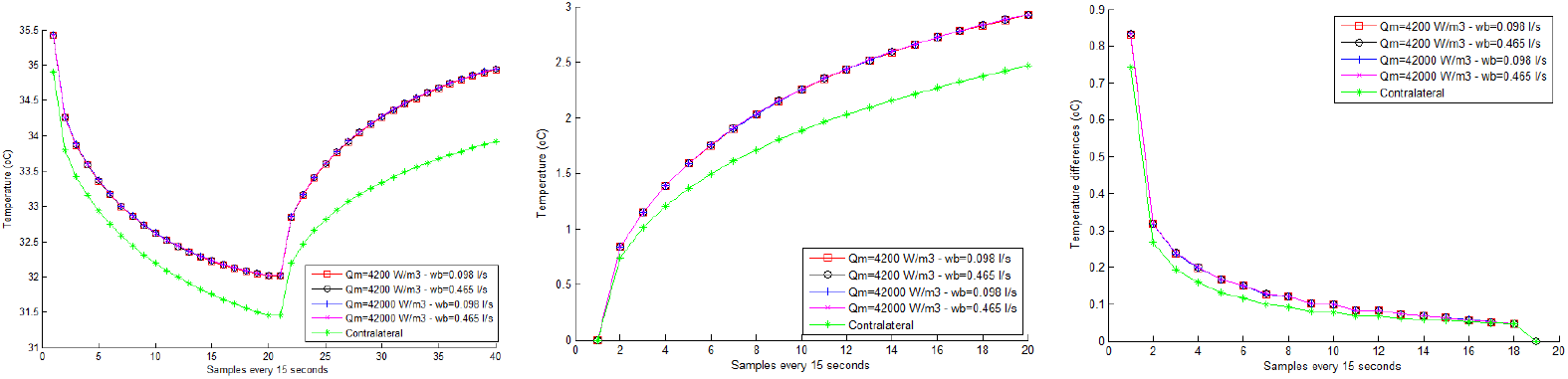
Influence of the metabolic heat and blood perfusion rate at transient state for possible combinations of nodular properties in front of it and on the symmetric neck position (on the red circle and cyan diamond of Fig. 5-left). Considering all simulation time (left). The reheating stage organizing considering all curves starting at the same temperature (center). The difference between consecutive temperatures, representing the heating rate (right).

However, it can be seen that there is a difference between the series of temperatures related to the placed on the skin in front of the nodule and its contralateral point (marked with the red circle and cyan diamond in Fig. 5-left). This means that it could be possible to distinguish if there is a nodule (at least for big nodules and skinny neck). Moreover, this difference is greater just after the end of the refrigeration process showing the importance of the examination under the induced change of temperature.

### Analysis of infrared examinations

This subsection shows the acquired temperatures in the exams of eight volunteers (patients on follow up in the university hospital) that are examined by using the protocol simulated by the numerical study. Several other details from previous exams, performed during their follow-up, are known (Table 2). The patients are selected when their exams and data better match the hypotheses considered in the numerical simulations.

A temperature series was organized from the infrared examinations of each patient. For it, firstly a point of the central nodule region (manually indicated by a doctor on the thermogram) was considered. Then, this location was used to compute the average temperature of an 11 × 11 window around it for each thermogram. The combination of them represents the patient. When two patients are compared their series are redrawn from the first value to show them starting at the same point, as shown in Fig. 11. The next sections present more details.

**Fig 11.**
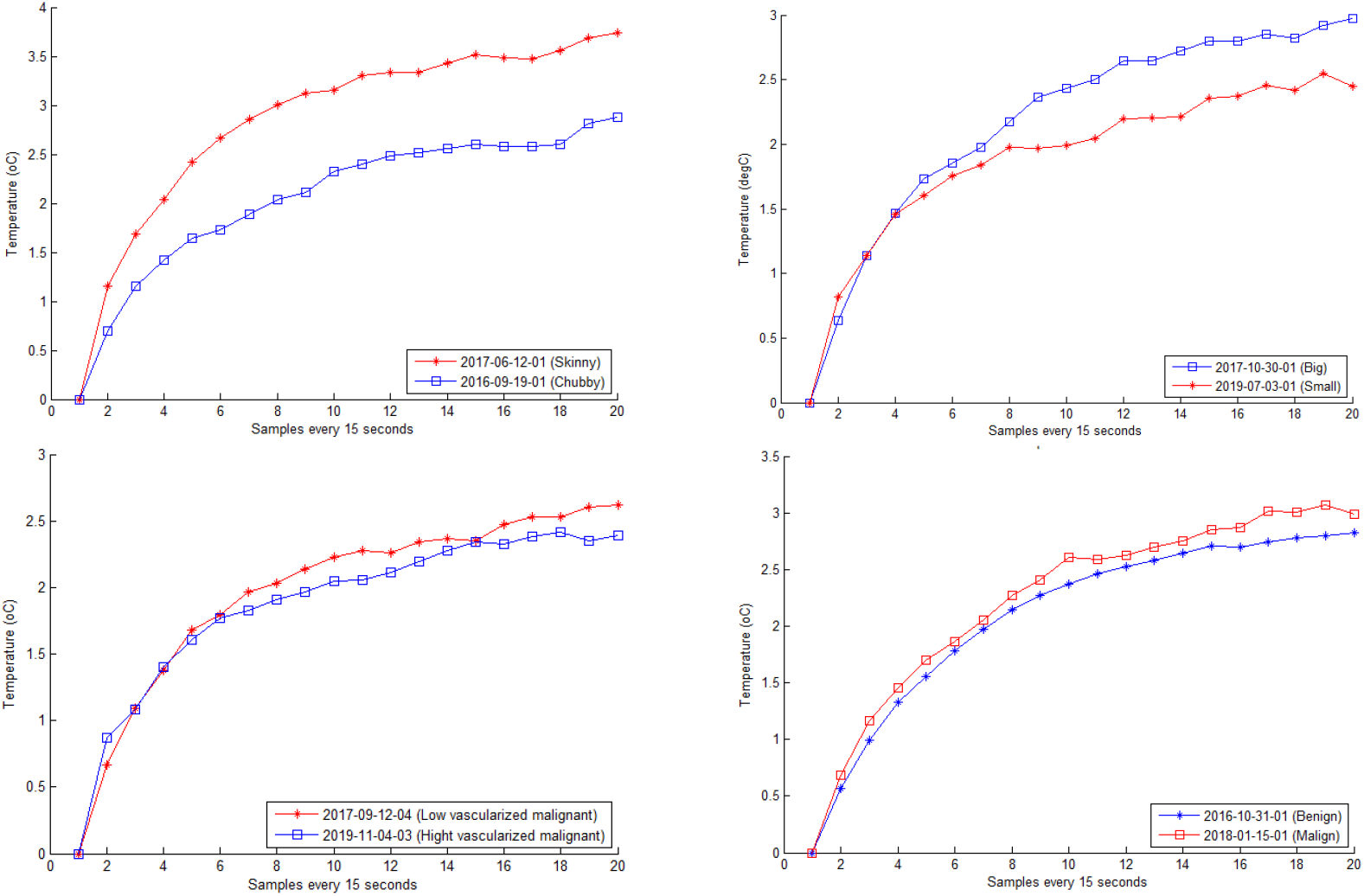
Temperature series in front of eight nodule of patients examined by an infrared camera. Each graph considers especial characteristics of nodules and patients numerically simulated.

#### Influence of fat tissue thickness

To analyze the influence of the fat tissue thickness on temperature series, the infrared examinations of the patient 2017-06-12-01 and 2016-09-19-01 were compared in this subsection (see them at Fig. 4 (a) and (b)). They have different body constitutions and fat tissue thickness. Patient 2017-06-12-01 (skinny) has less body mass index (BMI) than the patient 2016-09-19-01 (chubby). Patient 2017-06-12-01 has a unitary benign thyroid nodule. Patient 2016-09-19-01 has three benign nodules, the one more left in the (b) image of Fig. 4) was select for comparison. These nodules of both patients are similar considering their dimensions, ultrasonographic characteristics, and cytopathological result (Table 2).

Curves in Fig. 11-top-left show their temperature series (related to the same initial temperature). The series of the chubby patient (blue line) is below that of the skinny patient (red line) for the reheating stage. This shows the same behavior related to the thermal insulating effect of the fat tissue layer presented in the numerical modeling (Fig. 8-center).

#### Influence of nodule size

To analyze the influence of nodule size, we choose two patients presenting almost the same other characteristics except their nodular size. For this, the patient 2017-10-30-01 and 2019-07-03-01 were selected to have their series compared. These patients have a unitary malignant thyroid nodule with a similar pattern of vascularization and ultrasound characteristics (Table 2).

The nodule size of the patient 2017-10-30-01 was identified by ultrasound as being with diameters of (4.2 and 2.3 *cm*) or area = 30.4 *cm*^2^, that is around seven times bigger than the nodule of the patient 2017-10-30-01 (whose nodule was identified by ultrasound as being with diameters 1.4 and 1.0 *cm*, or with area = 4.4 *cm*^2^). Both patients have similar fat tissue thickness, and both nodules are located in the same thyroid area: the lobes. Fig. 4 (c) and (d) present them.

The curves in Fig. 11-top-right show the temperature in relation to the initial temperature of the patient. As can be seen, the series of the patient with the biggest nodule (blue line) has values greater than those of the patient with the smaller nodule (red line). It’s means that the bigger nodule has more influence on temperatures of skin surface than the smaller one, as shown in the numerical simulations (Fig. 9-center).

#### Influence of vascularization

As discussed earlier, the blood perfusion and the metabolic heat of nodules are important thermophysical parameters of the bioheat transfer theory analyzed in the previous simulations. Biologically, malignant nodules seem to be heat producers due to their cellular metabolism and characteristic of greater central vascularization than peripheral one. Although, there are reports of malignant nodules poorly vascularized. Some works [47, 48] report that there is not a well-defined relationship between the blood perfusion and the nodule vascularization.

Furthermore, if there are controversies regarding the relationship between malignancy and vascularization in the biological behavior of the tumor, the relationship between these and the usable parameters to model the behavior in simulations is even more complex, especially those related to the temperature that reaches the surface and can be detected by infrared sensors.

Two study related to the influence of vascularization are presented in this subsection. Firstly, two patients with malignant thyroid nodules but a low and a high vascularized nodules are compared. Then, the others two with a similar pattern of vascularization but presenting a malignant and benignant nodule are confronted.

The patients 2017-09-12-04 and 2019-11-04-03 were selected for the first comparison. The images in Fig. 4 (e) and (f) present them. They have a unitary malignant thyroid nodule with similar dimensions and different pattern of vascularization. Both are skinny, having a similar body mass index. The nodule of the patient 2017-09-12-04 has only peripheral vascularization (second level in the Chammas scale [50]) and the nodule of the patient 2019-11-04-03 has predominant central vascularization (fifth level in the Chammas scale) (Table 2).

Curves in Fig. 11-bottom-left show the temperatures concerning to the first value obtained from the infrared examinations. As can be seen, both series are very similar. This suggests that the difference of vascularization patterns of these nodules (with the same diagnostic, similar dimensions and ultrasonographic characteristics) causes little or no difference in the skin surface during the examination with infrared sensors. This fact was also observed in the numerical simulations (Fig. 10).

The patients 2016-10-31-01 and 2018-01-15-01 were selected for the second comparison. They have thyroid nodules with different diagnostic (Table 2) but nodules with similar dimensions and ultrasound characteristics, as shown in Table 2. Fig. 4 (g) and (h) present them. The patient 2016-10-31-01 has three benignant thyroid nodules, but only the biggest (the first from left to right in Fig. 4-g) was here compared. The patient 2018-01-15-01 has one malignant thyroid nodule. These nodules have peripheral vascularization (second and third levels in the Chammas scale [50] for the patients 2016-10-31-01 and 2018-01-15-01, respectively).

Curves in Fig. 11-bottom-right show the series concerning these nodules obtained from the infrared examinations. As can be seen, both are very similar. These two nodules with similar characteristics and similar patient’s characteristics cause a similar thermal effect, even with different diagnostic. This fact was also observed in the numerical simulations of Fig. 7. Then, the results of this subsection suggest that the thermal effects of thyroid nodules on the neck surface are not sufficient to differentiate benign from malign, and for this other features or methods must be considered.

## Discussion

The heat produced by thyroid nodules (malignant or benignant ones) could constitute visible patterns on the skin when acquired by infrared sensors under specific conditions [37, 51]. However, to understand the influence of all related parameters a numerical analysis must be done because, due to their diversity, each possible combination could not be easily found in human beings for identification of their influence on the skin temperature. Therefore, a parametric study is paramount to understand the influence of each element and to verify the potentiality of the thermography for diagnosis.

This work focuses on identifying how many parameters can do influence the heat transfer mechanism of biological tissues of the neck area and their relation to the detection of a thyroid nodule. For this, combinations of nodular sizes, fat tissue thicknesses, nodule blood perfusion rates, and nodule metabolic heat were analyzed. The protocol for infrared examination used in the University Hospital was the guide of all studies executed [42, 43]. Hereafter, a discussion about the principal results is presented.

Simulations at steady state show that there is a perceptible surface temperature variation between the point just in the front of the nodule and its contralateral point for a reduced fat tissue layer. The thermal influence of the nodules size and all other parameters on the skin surface decreases when the thickness of the fat layer increases. Almost no skin effect was detected for the larger fat layer (1.2 *cm*). Therefore, these simulations show that the thermal insulation effect of the fat tissue layer is much more important than the nodule size, although, both have the expected influence on surface temperature.

In the transient state, as well as in the steady state, the lower temperature variations were obtained in the skin surface when the major fat tissue thickness was considered, showing that this layer ever reduces the influence of the nodule in the neck surface. The larger nodule can be more easily detectable than the smaller nodules. No differences were observable for the lower nodule. To analyze the influence of metabolic heat and blood perfusion rate in the simulations, both steady state and transient state were evaluated and all combinations of these parameters were considered. For this, a large nodule (with diameters of 3.14 and 1.80 *cm*) was considered, and at the same time the adipose tissue was neglected (i.e. considered as 0 *cm* thickness layers, because the inclusion of such layer would only attenuate any difference, in case its existence). No differences were obtained for any combinations of the metabolic heat and blood perfusion rate performed. Therefore, whatever combination of them, within the range found in the literature, is the one representative of malignant or benign nodules, it is not possible to detect it on skin temperature. However, there is a difference between the temperature series of symmetric points (contralateral regions) on the surface of the skin when there is a nodule in one side when the nodule has enough size and there are few fat tissues. This means that there are conditions enabling to distinguish where there is a nodule considering the temperature in contralateral position, but disallow to recognize its classification (benign or carcinogenic).

Simulations at transient state also show that the final temperatures in the reheating stage are not the same that initial temperatures for all fat thickness considered. This probably shows that five minutes for the reheating stage is not enough to reach the body temperature before neck refrigeration (i. e. before the cooling stage).

Furthermore, temperatures of volunteers selected by patients on follow up in the university hospital, acquired by an infrared sensor (FLIR camera model SC620) using the simulated protocol [42, 43] were analyzed to verify the numerical findings. The temperatures on the neck surface of two patients differing by their body-mass index, but with nodules of similar dimensions, ultrasound characteristics, and the same diagnostic, show that the temperature changes were higher for the skinny patient as previewed by numerical modeling.

Other two patients differing as much as possible only by their nodule size, were considered to analyze this influence: temperatures of the patient with the larger nodule were greater than that with the small nodule, agreeing with the numerical simulations about the nodular size influence on skin temperatures. Beyond, two volunteers were selected to have their infrared examinations compared to the identification of the influence of vascularization. Their nodules had the same diagnostic, similar dimensions and ultrasound characteristics, and their thermography presented same behavior in the examination. This fact was also observed in the numerical simulations. Finally, the comparison of nodules of the two selected patients with similar characteristics but with different diagnostics, also shows little influence in the superficial temperatures. Confirming no relations among parameters of vascularization and degree of malignancy of the nodules.

## Conclusion

This work studies how several parameters of thyroid nodules (blood perfusion rate, metabolic heat, size, etc.) and the patient body characteristics (thickness of the fat tissue in the neck, for instance) can influence the identification of thyroid nodules by infrared sensors. For this, a bio-heat transfer analysis was performed through numerical simulations by using the COMSOL Multiphysics®v5.2 software for Finite Element modeling in order to identify possible relevant parameters. Moreover, infrared examinations of volunteers with nodule diagnostics were analyzed to verify if the numerical results are compatible with reality found in humans.

For the simulations, a 2D simplified geometry, based on a consensus image of an average neck, was created. The simulated neck considers skin, fat and muscle layers, thyroid gland, and an elliptic nodule inside its left lobe. The Pennes’ equation (in both transient and steady state) was used as mathematical and physical bioheat transfer model. Thermal convection (natural or forced) in the surface of the skin layer, thermal insulation in the trachea surrounding region, and prescribed temperature at 37 °*C* on the remaining surfaces, were considered as boundary conditions. Thermophysical parameters were investigated in the literature for each simulated tissue layer. The importance of thyroid nodule parameters (blood perfusion rate, metabolic heat, size) and body characteristics (thickness of the fat tissue in the neck) were analyzed at transient and steady states. Four fat tissue thickness, three different dimensions of the nodule, and four combinations of blood perfusion and metabolic heat of the nodule were considered in simulation in order to analyze their influence on possible nodule diagnosis.

The simulations show the great thermal insulation effect of the fat tissue layer. This means that the influence of the heat produced by a thyroid nodule on the surface of the skin will decrease if the layer of fatty tissue is large. Inversely, simulations with any combination of metabolic heat and blood perfusion rate of the thyroid nodules could not promote additional difference in temperature on the skin surface.

The analysis of the patient’s infrared examinations agrees with all findings. They show that the fat layer thickness presents great importance on thermal insulating effect. Moreover, differences in vascular patterns of nodules (with the same diagnostic, similar dimensions, and ultrasonographic characteristics) have no influence on the temperature of the surface during the examination of real patients confirming the results of the numeric study. Considering that vascularization could be related to the malignity of the nodules in the thyroid, the results indicate difficulties for identification of nodular malignancy by thermography considering equilibrium with the environment of even forced ventilation, although it is possible to identified a thyroidal nodule by using thermography.

### Future works

For thermography, an immediate consequence of this works is related to the importance of estimating the fat layer in the neck due to its relevance in the analysis and possible association with unfair diagnosis of nodules by IR. The amount of fat in the neck can be evaluated directly by using US or at least estimated considering the neck perimeter and the BMI of the patients. Moreover, it is relevant to investigate other acquisition protocols to analyze the infrared examinations in order to present final arguments related to their use (or do not) for cancer diagnosis of thyroid gland [42], [43].

For the heat simulation aspect, the relation between thermophysical parameters and the biological behavior must be better investigated. They can be measured experimentally in an approach similar to that used for the other tissues of the human body, or they can be estimated through the inverse problem method with thermal examinations of real patients, as well [52]. After, but yet related to simulation: (1) more generic elements and tissue surrounding the neck and their properties (like veins and trachea ventilation) could be included; (2) real geometry of an specific case based, for instance, on magnetic resonance or computer tomography can be modeled; and (3) more nodular elements (shape, size, 3D positions, heterogeneity, and structural constitution) can be included to help in confirming if the diagnosis of malignancy of thyroid nodules is really impossible (or possible in what conditions) by using this type of examination. These could then finally answer any doubt in this line of the research that begging in [28] and to what this work promote new conclusions simulating the transient behavior.

## Supporting information

**S1 File. Simulated geometry**. The simulated geometry in the geometry.mphbin file. This geometry was constructed in COMSOL Multiphysics Software v5.2.

**S2 File. Infrared examination file**. This 2016-09-19-01.rar file contains the infrared examination of the patient 2016-09-19-01.

**S2 File. Infrared examination file**. This 2016-10-31-01.rar file contains the infrared examination of the patient 2016-10-31-01.

**S2 File. Infrared examination file**. This 2017-06-12-01.rar file contains the infrared examination of the patient 2017-06-12-01.

**S2 File. Infrared examination file**. This 2017-09-12-04.rar file contains the infrared examination of the patient 2017-09-12-04.

**S2 File. Infrared examination file**. This 2017-10-30-01.rar file contains the infrared examination of the patient 2017-10-30-01.

**S2 File. Infrared examination file**. This 2018-01-15-01.rar file contains the infrared examination of the patient 2018-01-15-01.

**S2 File. Infrared examination file**. This 2019-07-03-01.rar file contains the infrared examination of the patient 2019-07-03-01.

**S2 File. Infrared examination file**. This 2019-11-04-03.rar file contains the infrared examination of the patient 2019-11-04-03.

All these examinations were performed using the FLIR infrared camera model SC620 and the protocol cited in [30]. These examinations are also available in http://visual.ic.uff.br/en/thyroid/.

## Acknowledgments

J.R.G.M. and M.B.H.M. are supported by the CAPES Brazilian Foundation. A.C. is partially supported by MACC-INCT, CNPq Brazilian Agency (402988/2016-7, and 305416/2018-9) and FAPERJ (project Tematicos: 210.019/2020).

